# A mechanical memory of pancreatic cancer cells

**DOI:** 10.1101/730960

**Authors:** Ilaria Carnevale, Mjriam Capula, Elisa Giovannetti, Thomas Schmidt, Stefano Coppola

## Abstract

Cells sense and respond to mechanical stimuli in healthy and pathological conditions. Although the major mechanisms underlying cellular mechanotransduction have been described, it remains largely unclear how cells store information on past mechanical cues over time. Such mechanical memory is extremely relevant in the onset of metastasis in which cancer cells migrate through tissues of different stiffness, e.g. from a stiffer tumor microenvironment to softer metastatic sites as commonly occurs for pancreatic cancer. Here, we used micropillar-based traction force microscopy to show that Suit-2.28 pancreatic cancer cells mechanically primed on a stiff matrix exerted higher traction forces even when transferred to a soft secondary matrix, as compared to soft-primed cells. This mechanical memory effect was mediated by the Yes-associated protein (YAP) and the microRNA-21 (miR-21) that are two mechanosensors initially identified as long-term memory keepers in mesenchymal stem cells. Soft-primed cells showed (i) a lower YAP nuclear translocation when transferred to a stiff secondary matrix and (ii) a loss of rigidity sensing through YAP, as compared to stiff-primed cells. The mechanical adaptation resulted in a differential expression of miR-21, inversely proportional to the priming rigidity. The long-term mechanical memory retained by miR-21 unveiled a previously unidentified mechanical modulation of drug resistance by past matrix stiffness. The higher expression of miR-21 in soft-primed cells correlated with the increased resistance to gemcitabine, as compared to stiff-primed and non-primed pancreatic cancer cells.

## Introduction

Cells are exposed to several mechanical and physical cues within their three-dimensional microenvironment, to which they respond by exerting forces, regulating their shape, internal cytoskeletal tension, and elastic modulus (1–5). These complex mechanoresponses are the result of integrated signals from many distinct mechanosensitive structures such as ion channels, focal adhesions, adherens junctions, cytoskeleton, and LINC complex (6, 7). Although the major mechanisms underlying cellular mechanotransduction have been clarified, the intriguing ability of cells to store information on past physical stimuli over time for future mechanical challenges remains largely unexplored (8).

The long-term mechanical memory of cells is predominantly mediated by changes in gene expression through the nuclear localization of transcription factors that act as memory keepers for days (9–12). Yang et al. demonstrated that for human mesenchymal stem cells, after a prolonged culture on stiff substrates, the Yes-associated protein (YAP) re-localization into the cytosol was prevented for up to 10 days when cells were transferred to softer environments (9). Such YAP-mediated memory was found to regulate the collective migration of mechanically primed epithelial cells (12). Li et al. identified the microRNA miR-21 as a memory keeper of the fibrogenic program in mesenchymal stem cells (11). During stiff priming, the nuclear translocation of the mechanosensitive myocardin-related transcription factor-A (MRTF-A) directly controlled the transcription of miR-21, whose levels remained high for 4-6 days after removal of the mechanical stimulus.

In this work, we show evidence that soft- and stiff-primed Suit-2.28 pancreatic cancer cell lines differentially exerted forces depending on past matrix stiffness (i.e. higher priming rigidity, larger forces on secondary stiffness), as quantified by micropillar-based traction force microscopy. Such mechanical memory effect was mediated by YAP activity (i.e. nuclear translocation) and miR-21 expression. The long-term memory of miR-21 elucidated the role of mechanical priming in the modulation of chemoresistance by showing that soft-primed pancreatic cancer cells were less sensitive to gemcitabine treatment, as compared to stiff-primed and non-primed control cells.

## Results and Discussion

We cultured Suit-2.28 cells, a pancreatic cancer cell line derived from a metastatic liver tumor (14), on fibronectin-coated soft (Young’s modulus E = 10 kPa) and stiff (E = 100 kPa) polydimethylsiloxane (PDMS) gels casted in multi-well plates for up to four passages, with 3 days/passage (Fig. 1A and the Supplementary Material). After the mechanical priming (i.e. P4), soft- and stiff-primed Suit-2.28 cells were transferred to (i) PDMS micropillars (effective Young’s modulus E = 9.8-137.1 kPa) to quantify cellular forces (Fig. 1B), (ii) PDMS gels (E = 10-100 kPa) to assess the YAP nuclear translocation (i.e. activity) (Fig. 1C), and (iii) plastic multiwell plates to evaluate the cell growth inhibition by gemcitabine (Fig. 1D). The secondary stiffness was either kept the same or switched compared to the priming stiffness.

**Fig. 1.**
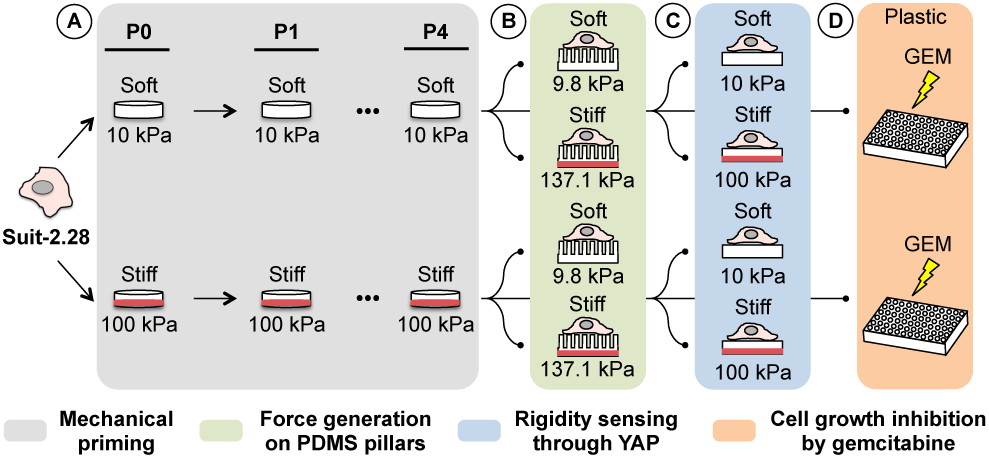
Experimental design. (A) Mechanical priming of Suit-2.28 pancreatic cancer cells. Cells were cultured on fibronectin-coated soft (Young’s modulus E = 10 kPa) and stiff (E = 100 kPa) PDMS substrates for four passages (P) every 3 days to mechanically adapt them. (B) After P4, soft- and stiff-primed cells were transferred to fibronection-coated soft (effective E = 9.8 kPa) and stiff (effective E = 137.1 kPa) elastic PDMS micropillars to quantify the effect of the mechanical priming on cellular forces (13). (C) The rigidity sensing through YAP activity was measured by transferring soft- and stiff-primed cells (after P4) to secondary stiffness (soft and stiff) and quantifying the nuclear translocation of YAP. (D) Cell growth inhibition by gemcitabine (GEM) at increasing concentration (10-1000 nM) was evaluated for soft- and stiff-primed cells transferred to plastic plates.

To assess the effect of the mechanical priming on the force machinery, we employed elastic micropillars whose deflections report on cellular forces (13) (Fig. 1B and Fig. 2A). As previously described for other cell lines (15), primed Suit-2.28 cells responded to changes in substrate rigidity by significantly increasing their size from soft (E = 9.8 kPa) to stiff (E = 137.1 kPa) pillars, i.e. from a median cell size of 327 *µm*^2^ to 384 *µm*^2^ for ‘Soft *→* Soft’ and ‘Soft *→* respectively and from 329 *µm*^2^ to 365 *µm*^2^ for ‘Stiff *→* Soft’ and ‘Stiff *→*Stiff’ respectively (Fig. 2B). Remarkably, the average force per pillar exerted by primed Suit-2.28 cells showed a previously unidentified mechanical memory effect (Fig. 2C). Stiff-primed cells always applied larger forces compared to soft-primed cells on both soft (from 2.0 *±* 0.7 nN to 2.5 *±*0.9 nN for ‘Soft *→* Soft’ vs.. ‘Stiff *→* Soft’, respectively) and stiff (from 24 *±*7 nN to 26 *±* 9 nN for ‘Soft *→*Stiff’ vs. ‘Stiff *→* Stiff’, respectively) micropillars (Fig. 2C), although no significant difference in cell spreading area was observed (same conditions as in Fig. 2B).

**Fig. 2.**
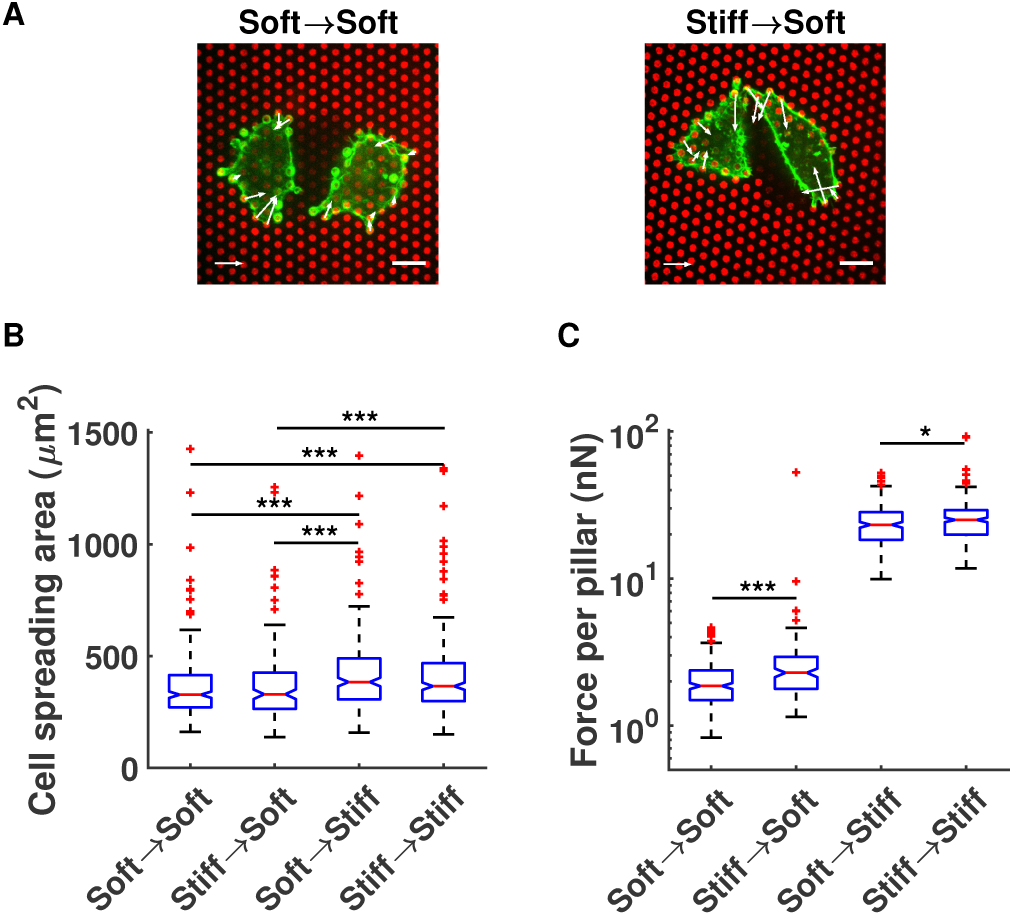
Cellular forces depend on the priming stiffness. (A) Representative confocal images of soft- and stiff-primed Suit-2.28 cells (actin in green) on soft PDMS micropillars (fibronectin-coated pillar tops in red). Scale bar: 10 *µm*. Force bar: 3 nN. (B) Cell spreading area of soft- and stiff-primed Suit-2.28 cells transferred to soft and stiff fibronectin-coated PDMS micropillars (e.g. ‘Soft *→* Soft’ refers to soft-primed cells transferred to a soft secondary stiffness). (C) Average force per pillar exerted by soft- and stiff-primed Suit-2.28 cells transferred to soft and stiff micropillars. The boxplot shows the median (central mark) and the 25th and 75th percentiles (bottom and top edges, respectively). N = 247, 138, 222, 265 cells for the four conditions, respectively, pooled together from three independent micropillar arrays (i.e. three independent experiments). *, *** indicate p<0.05 and p<0.001 significance levels for a Mann-Whitney U test.

Previous studies identified YAP (9, 12) and miR-21 (11) as effectors of mechanical memory. To address whether the observed memory effect in force generation was mediated by such mechanosensors, we investigated the activity of YAP (Fig. 1C and Fig. 3A) and the expression of miR-21 in primed Suit-2.28 cells (Fig. 1D).

**Fig. 3.**
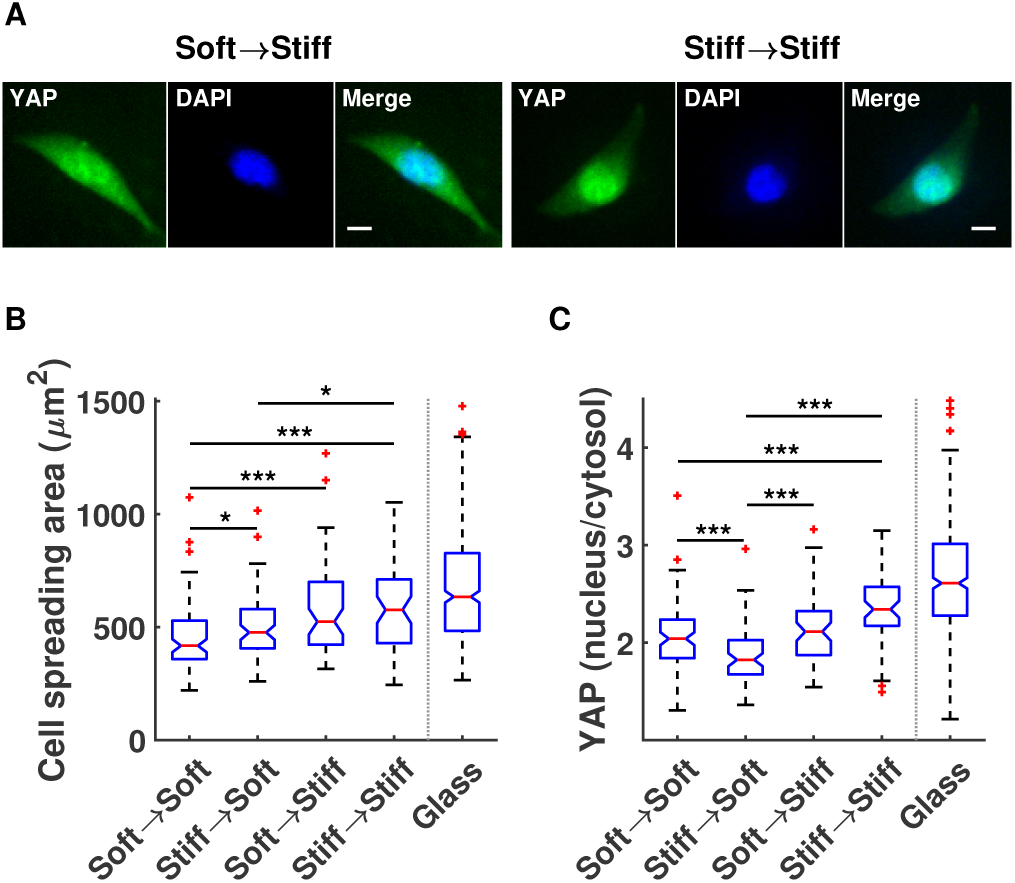
YAP stores mechanical memory. (A) Representative confocal images of the YAP signal (green) in soft- and stiff-primed Suit-2.28 cells on stiff PDMS sub-strates. DAPI (blue) was used to locate and segment the nucleus. Scale bar: 10 *µm*. (B) Cell spreading area of soft- and stiff-primed Suit-2.28 cells transferred to soft and stiff fibronectin-coated PDMS substrates (e.g. ‘Soft *→* Soft’ refers to soft-primed cells transferred to a soft secondary stiffness). (C) YAP activity was measured by quantifying the nuclear-to-cytosol ratio of the YAP fluorescent intensity for soft- and stiff-primed Suit-2.28 cells transferred to soft and stiff substrates. The ‘Glass’ condition is used as a reference and indicates non-primed cells seeded on a fibronectin-coated glass substrate. The boxplot shows the median (central mark) and the 25th and 75th percentiles (bottom and top edges, respectively). N = 87, 70, 58, 55, 200 cells for the five conditions, respectively, pooled together from two independent PDMS substrates (i.e. two independent experiments). *, *** indicate p<0.05 and p<0.001 significance levels for a Mann-Whitney U test, only within the mechanical priming conditions.

While the past matrix stiffness affected the cell size only when primed cells were transferred to a soft secondary stiffness (median cell size from 418 *µm*^2^ to 476 *µm*^2^ for ‘Soft *→*Soft’ and ‘Stiff *→* Soft’ respectively in Fig. 3B), YAP stored the mechanical memory (Fig. 3C). The average nuclear-to-cytosol ratio of YAP fluorescence intensity significantly decreased from 2.1 *±*0.3 to 1.9 *±* 0.3 when soft- and stiff-primed cells adhered on the soft substrate respectively, whereas it significantly increased from 2.1 *±*0.4 to 2.4 *±* 0.4 when primed Suit-2.28 cells were seeded on the stiff substrate (3C). It is interesting to note that the rigidity sensing through YAP was lost for soft-primed cells (’Soft *→*Soft’ vs. ‘Soft *→* Stiff’ in Fig. 3C), whereas retained for stiff-primed cells (’Stiff *→*Soft’ vs. ‘Stiff *→* Stiff’ in Fig. 3C). The expected increase in cell size from soft to stiff substrates (median cell size from 418 *µm*^2^ to 524 *µm*^2^ for ‘Soft*→*Soft’ vs. ‘Soft *→* Stiff’ in Fig. 3B) suggests that for soft-primed Suit-2.28 cells the rigidity sensing was potentially mediated by other mechanosensors than YAP, e.g. focal adhesions as reported in (12). As controls, we also report the significantly higher cell spreading area and YAP nuclear localization of non-primed Suit-2.28 cells adherent on fibronectin-coated glass (E ∼ 1 GPa) (Fig. 3B-C). Our findings on YAP’s ability to store the mechanical memory show a different and to some extent more complex response to the mechanical adaptation with respect to the work by Nasrollahi et al. (12) in which stiff-primed cells always showed higher YAP nuclear localization compared to soft-primed cells, independently of the secondary stiffness. Such different picture may be caused by e.g. stiffness ranges (0.5 kPa vs. 50 kPa in (12) compared to our 10 kPa vs. 100 kPa), priming timescales, cell types, and cell confluency (i.e. a monolayer vs. single cell confluency highly influence YAP nuclear translocation (16)) as confounding factors.

The expression of miR-21 significantly changed over the course of the mechanical priming as quantified by qRT-PCR (Fig. 4A). At low passage number there was no significant difference of the relative miR-21 gene expression between (soft- and -stiff) primed and non-primed (’Plastic’) Suit-2.28 cells (P1, Fig. 4A), whereas at the latest passage (after ∼9 days of mechanical priming) a 3 to 3.8-fold increase was observed in stiff- and soft-primed cells, respectively (P4, Fig. 4A). This finding contributes to an increasing body of evidence on mechanosensitive miRNAs which mediate cellular responses to mechanical stimuli like shear stress (17) and matrix strain (11, 18). It is also interesting to note that compared to mesenchymal stem cells (11), we observed an upregulation of miR-21 in soft vs. stiff substrates for pancreatic cancer cells. Previous studies have shown a correlation between high miR-21 expression and increased resistance to gemcitabine (19), a nucleoside analog drug used as a first-line treatment in pancreatic cancer (20). To test whether the observed differential expression of miR-21 in soft-vs. stiff-primed cells could modulate the sensitivity to gemcitabine, we performed a sulforhodamine B (SRB, (21)) growth inhibition assay (Fig. 1D). For such drug assay, we decided to transfer all primed cells (i.e. after P4, Fig. 1A) to plastic multiwell culture plates and expose them to gemcitabine for 36 hours with the aim to further testing the long-term mechanical memory of miR-21, that was estimated to keep it for 4-6 days in mesenchymal stem cells (11). At the highest drug concentrations (i.e. 100 and 1000 nM), soft- and stiff-primed cells showed a 2 to 50-fold significant reduction in sensitivity to gemcitabine compared to non-primed Suit-2.28 cells (Fig. 4B). Remarkably, the response to gemcitabine as a function of priming stiffness was scaling comparably to the relative miR-21 expression (Fig. 4A). At the lowest concentration (i.e. 10 nM), although the stiff-primed cells died significantly more than non-primed ones, the growth of soft-primed cells was significantly less inhibited by gemcitabine (Fig. 4B). To the best of our knowledge, these findings demonstrate for the first time the role of the mechanical memory in modulating the sensitivity of pancreatic cancer cells to gemcitabine through the regulation of miR-21 expression. A recent work by Rice et al. (22) reported that stiffness induces chemoresistance to paclitaxel, but not to gemcitabine in pancreatic cancer cells. However, differently to our approach, the drug assay in (22) was performed by seeding cells on gels for 72 hours without any prior and long-term mechanical priming, i.e. the effect of the mechanical adaptation on resistance regulators such as miR-21 was potentially not still significant (as in P1, Fig. 4A).

**Fig. 4.**
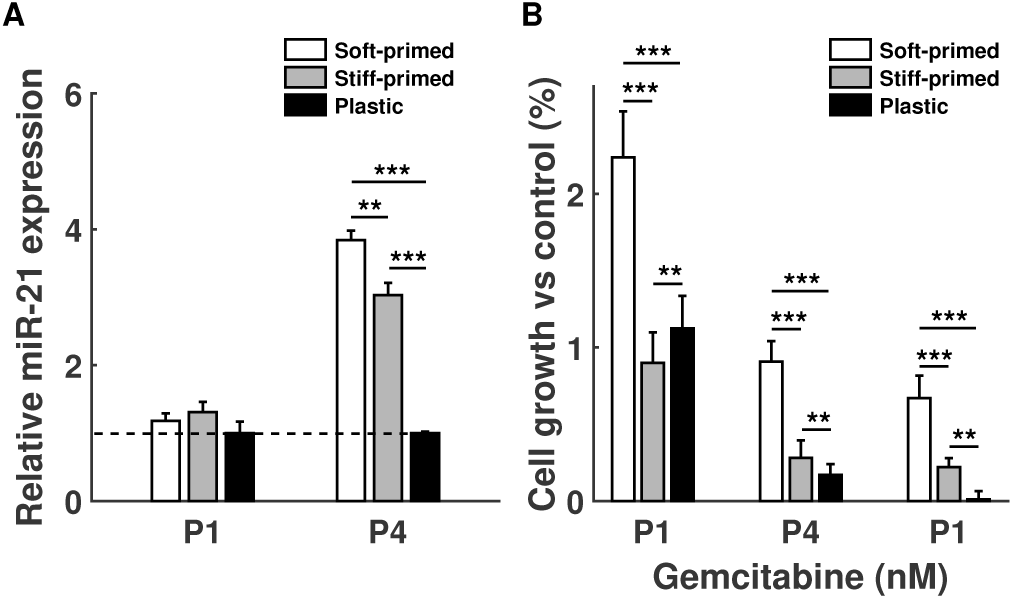
Gemcitabine sensitivity depends on the priming stiffness through miR-21 expression. (A) Relative miR-21 gene expression of soft-, stiff-primed and non-primed Suit-2.28 cells at passage P1 and P4 (see Fig. 1A) as quantified by qRT–PCR. Expression levels of miRNAs were first normalized to housekeeping RNU6 in all experimental conditions and then expressed as relative levels to Plastic at passage P1 and P4, respectively. Values represent mean*±*SD of three separate tests. **, *** indicate p<0.01 and p<0.001 significance levels for a t-test, performed within the same passage. (B) Evaluation of cell growth inhibition of gemcitabine at increasing concentrations (i.e. 10, 100, 1000 nM for 36 hours exposure) for soft-, stiff-primed and non-primed Suit-2.28 cells (’Soft-primed’, ‘Stiff-primed’, and ‘Plastic’, respectively) at passage P4 (see Fig. 1D), as quantified by sulforhodamine B (SRB) assay (21). Values represent mean*±*SD of six separate tests. **, *** indicate p<0.01 and p<0.001 significance levels for a t-test, performed within the same drug concentration.

## Conclusions

Overall, the picture presented here demonstrates that Suit-2.28 pancreatic cancer cells show a mechanical memory effect through YAP and miR-21 as memory keepers, when adapted to substrates of varying stiffness. While weakly regulating cell size, such mechanical memory significantly affected the forces generated by cells in response to the change in substrate rigidity. The decreased sensitivity to gemcitabine observed in soft-primed cells suggests that the environmental rigidity plays a role in modulating the gemcitabine chemoresistance in pancreatic cancer. Notably, while pancreatic cancer cells grow within a dense, stiff stroma at their primary site, the metastatic niche in the liver (the major metastatic site) is characterized by a softer microenvironment. Yet, liver metastases present a high intrinsic resistance, e.g. the radiological response rate to gemcitabine monotherapy at both primary and metastatic sites is below 10% (23, 24). In addition to the genetic heterogeneity of metastasis (25), these findings are potentially supported by our results on how pancreatic cancer cells acquire a more aggressive/resistant phenotype after a long-term mechanical adaptation to a softer microenvironment.

Further studies are required to unravel whether the molecular pathways related to memory are potentially modulated by matrix composition (i.e. the role of integrin signaling) in addition to past matrix stiffness.

## Experimental procedures

### Cell culture and mechanical priming

Human pancreatic ductal adenocarcinoma Suit-2.28 cells (14) were cultured using RPMI 1640 medium (Lonza Walkersville Inc., Walkersville, MD, USA) containing fetal bovine serum (FBS, 10%), penicillin (1%) and streptomycin (1%) and cells have been kept in a CO_2_ incubator (NuAire, Plymouth, MN, USA), at 5% CO_2_ and 37*°*C.

The mechanical priming was performed by culturing for several passages (P0-P4, every 3 days) Suit-2.28 cells (100,000 cells/well) in plastic six-well plates (Sarstedt, Numbrecht, Germany) in which polydimethylsiloxane (PDMS) (Sylgard 184, Dow Corning, Midland, MI, USA) gels were cast. For every passage, four independent wells were used (i.e. four independent mechanical priming experiments were performed). PDMS was mixed with 1:30 or 1:60 crosslinker:base ratios, degassed, poured into the plates, and cured for 4 hours at 80*°*C. As described in (26), such curing time and temperature yield PDMS gels with a Young’s modulus E of ∼100 kPa (referred as ‘Stiff’ substrates in the manuscript) and ∼10 kPa (referred as ‘Soft’ substrates in the manuscript), for 1:30 and 1:60 crosslinker:base ratios, respectively. All gels used in the study were produced (i.e. PDMS mixing and curing) at the same time to reduce variability in substrate stiffness. Prior to cell seeding, each PDMS gel-containing well was coated with fibronectin (F1141-1MG, Sigma-Aldrich Chemie N.V., Zwijndrecht, The Netherlands) at a final concentration of 2 *µ*g/cm^2^ for 1-2 hours at 37*°*C, after surface activation by UV-ozone cleaner (Jelight, Irvine, CA, USA) for 10 minutes.

### Micropillar-based traction force microscopy

Cellular traction force measurements were performed using elastic micropillar arrays produced in our labs. A hexagonal array of poly-di-methyl-siloxane (PDMS, Sylgard 184, Dow Corning, Midland, MI, USA) micropillars of 2 *µ*m diameter, 4 *µ*m center-to-center distance and with a height of 6.9 *µ*m (Young’s modulus 9.8 kPa effective stiffness) or 3.2 *µ*m (137.1 kPa effective stiffness) were produced using replica-molding from a silicon wafer (13). The pillar arrays were flanked by integrated 50 *µ*m high spacers to allow the inversion onto glass coverslips, without compromising the limited working distance of a high-NA objective on an inverted microscope. The tops of the micropillars were coated with a mixture of unlabeled (F1141-1MG, Sigma-Aldrich Chemie N.V., Zwijndrecht, The Netherlands) and Alexa Fluor 647-labeled fibronectin (1:5) using micro-contact printing. The position of the pillar tops was observed by confocal fluorescence microscopy and determined down to sub-wavelength accuracy using custom software (Matlab, Mathworks, Natick, MA, USA). Forces were obtained by multiplying the pillar deflections by the array’s characteristic spring constant (13.7 nN/*µ*m and 191.4 nN/*µ*m, respectively, determined by finite element modeling) (13). The pillars’ spring constants were converted to an equivalent Young’s modulus for continuous substrates (27) of 9.8 kPa (soft pillars) and 137.1 kPa (stiff pillars), respectively. Only pillars closer to the cell perimeter than 3 *µ*m and with a deflection *>*68 nm for soft pillars and *>*76 nm for stiff pillars were considered for the calculation of the average force per pillar and the total cellular forces. The deflection thresholds, which reflect the positional accuracy by which individual pillars could be localized, were determined for each confocal image as the 75th percentile of the displacements of pillars outside the cell area (i.e. not bent by the cells). The cell spreading area was determined by thresholding the fluorescence signal of Alexa Fluor 532-Phalloidin (Thermo Fisher Scientific, Waltham, MA, USA) labeled actin filaments using a triangular threshold method (28).

### Immunofluorescence and confocal microscopy

Six-teen hours after seeding on the secondary stiffness, cells were fixed in 4% Paraformaldehyde (PFA) at room temperature (RT) for 15 minutes. After washing with 1X PBS, cells were permeabilized with 0.1% Triton X-100 for 10 minutes at RT, washed with 1X PBS and incubated with 3% bovine serum albumin (BSA) at RT for at least 1 hour. Cells for the YAP activity measurements were first incubated with primary YAP antibody (1:50, sc101199, Santa Cruz Biotechnology, Dallas, TX, USA) overnight at 4*°*C and after PBS washing were incubated with DAPI (1:1000, Thermo Fisher Scientific, Waltham, MA, USA) and Alexa Fluor 532 goat anti mouse (1:200, Thermo Fisher Scientific, Waltham, MA, USA) for 1 hour at RT. Cells on micropillars were directly incubated with Alexa Fluor 532 Phalloidin (1:1000, Thermo Fisher Scientific, Waltham, MA, USA) for 1 hour at RT.

Imaging was performed on a home-built setup based on an Axiovert200 microscope body (Zeiss, Oberkochen, Germany). Confocal imaging was achieved by means of a spinning disk unit (CSU-X1, Yokogawa, Tokyo, Japan). The confocal image was acquired on an emCCD camera (iXon 897, Andor, Oxford Instruments, Abingdon, UK). IQ-software (Andor, Oxford Instruments, Abingdon, UK) was used for basic setup-control and data acquisition. Illumination of DAPI, Alexa Fluor 532-Phalloidin, Alexa Fluor 532 goat anti-mouse, and Alexa Fluor 647-fibronectin was performed with three different lasers of wavelength 405 nm (CrystaLaser, Reno, NV, USA), 514 nm (Cobolt, Solna, Sweden), and 642 nm (Newport Spectra-Physics BV, Utrecht, The Netherlands). Accurately controlled excitation intensity and excitation timing were achieved using an acousto-optic tunable filter (AA Optoelectronics, Toronto, Canada). Light was coupled into the confocal spinning-disk unit by means of a polarization maintaining single-mode fiber (OZ Optics, Ottawa, Canada). The fluorescent signal was collected by a 20X/0.5 air objective (Zeiss, Oberkochen, Germany) for YAP activity measurements or 100X/1.4 oil objective (Zeiss, Oberkochen, Germany) for micropillar-based traction force microscopy.

### YAP activity

To quantify the nuclear translocation of YAP (i.e. YAP activity), the mean fluorescence intensity of YAP was measured in the nucleus and the cytoplasm and their ratio was calculated. The DAPI signal was used to locate and segment the nuclei. The cell spreading area was determined by thresholding the fluorescence signal of Alexa Fluor 532 labeled YAP using a triangular threshold method (28). All image analysis was performed by custom software (Matlab, Mathworks, Natick, MA, USA).

### Quantitative real-time PCR

Cell pellets were collected during the priming study at passages P1 and P4. Cells were harvested in a tube with 250 *µ*l of Trizol reagent (Sigma-Aldrich Chemie N.V., Zwijndrecht, The Netherlands). Total mRNA has been extracted by adding 50 *µ*l of chloroform to each sample, shaking vigorously and spinning for 15 minutes at 12,000 *g* at 4*°*C, after 5 minutes of incubation at room temperature. To allow the precipitation of mRNA, the aqueous phase has been washed with 125 *µ*l isopropanol and 500 *µ*l of 70% ethanol. The pellet has been finally resuspended in nuclease free water and the amount of nucleic acid determined through NanoDrop technology (ND 1000, Thermo Fisher Scientific, Waltham, MA, USA). Ten ng of total RNA was subjected to reverse-transcription with the miRCURY LNA miRNA PCR System (QIAGEN, Venlo, The Netherlands), following the manufacturer’s instructions. The expression levels of miR-21 were quantified by qRT-PCR using miRCURY SYBR Green PCR Kit (QIAGEN, Venlo, The Netherlands). The results, normalized to the RNU6 house-keeping gene as comprehensively investigated in (29) for several pancreatic cancer cell lines, are represented as the fold expression *±* standard deviation (SD) with respect to the non-primed condition (i.e. ‘Plastic’).

### SRB assay

Cell growth inhibition was performed with the sulforhodamine B assay (SRB), as previously described in (21). Briefly, cells in exponential growth phase were harvested by trypsinization, counted with the Coulter Counter (Z Series, Beckman, Indianapolis, USA) and plated at concentrations of 3000 cells per well in a 96-well flat bottom plate (Greiner Bio-One GmbH, Frickenhausen, Germany). Eight hours after plating, cells were treated with gemcitabine at increasing concentrations (i.e. 10, 100, 1000 nM). After 36 h of treatment, cells were fixed with 50% TCA (25*µ*l/well) at 4*°*C during 1 h and washed 5 times with Milli-Q H_2_O. Next, the cells were stained for 15 min with 0.4% SRB (Sigma-Aldrich, Zwijndrecht, Netherlands) in 1% acetic acid and washed 4 times with 1% acetic acid to remove the unbound stain. Plates were allowed to dry, after which the protein-bound stain was dissolved in 150 *µ*l of 10mM Tris-base (Sigma-Aldrich, Zwijndrecht, Netherlands). The optical density was read at 540 nm with the HTX Synergy microplate reader (BioTek, Munich, Germany).

## Author Contributions

S.C. and I.C. designed the experiments. S.C., I.C., and M.C. performed the experiments. S.C. analyzed the data. E.G. and T.S. supervised the research. S.C. and I.C wrote the article.

## Acknowledgements

We thank Btissame El Hassouni for helping with cell culture and the drug inhibition assay. S.C. was beneficiary of an AXA Research Fund Post-doctoral grant. I.C. was beneficiary of a postdoctoral fellowship by Associazione Medica Forlivese “Giovanni Fontana” - Associazione Italiana per lo Studio del Pancreas (AISP). This work was partially supported for the materials and collection/analysis of data by the KWF Dutch Cancer Society grant (KWF project #10401, EG) and Cancer Center Amsterdam (CCA-2015-1-19 grant, EG).

